# Temporal differentiation of bovine airway epithelial cells grown at an air-liquid interface

**DOI:** 10.1101/197814

**Authors:** Daniel Cozens, Erin Sutherland, Francesco Marchesi, Geraldine Taylor, Catherine Berry, Robert Davies

**Affiliations:** Institute of Infection, Immunity and Inflammation, College of Medical, Veterinary and Life Sciences, University of Glasgow, Glasgow, UK; School of Veterinary Medicine, University of Glasgow, Glasgow, UK; The Pirbright Institute, Pirbright, Surrey, UK; Institute of Molecular, Cell and Systems Biology, College of Medical, Veterinary and Life Sciences, University of Glasgow, Glasgow, UK

## Abstract

The respiratory epithelium is exposed to assault by toxins and pathogens through the process of inhalation, which has numerous implications on both human and animal health. As such, there is a need to develop and characterise an *in vitro* model of the airway epithelium to study respiratory pathologies during infection or toxicology experiments. This has been achieved by growing airway epithelial cells at an air-liquid interface (ALI). Characterisation of ALI models are not well-defined for airway epithelial cells derived from non-human species. In this study we have fully characterised a bovine airway epithelial cell models (AECM) grown at an ALI in relation to *ex vivo* tissue. The morphology of the model was monitored at three day intervals, to identify the time-period at which the culture was optimally differentiated. The model was shown to be fully-differentiated by day 21 post-ALI. The culture formed a stereotypical pseudostratified, columnar epithelium containing the major cell types of the bronchial epithelium (ciliated-, goblet- and basal cells). Once fully differentiated the bovine AECM displayed both barrier function, through the formation of tight-junctions, and active mucociliary clearance, important properties of the mucosal barrier. The bovine bronchial epithelial cells remained stable for three weeks, with no evidence of deterioration or dedifferentiation. The window in which the model displayed full differentiation was determined to be between day 21-42 post-ALI. Through comparison with *ex vivo* tissue derived from donor animals, our bovine AECM was shown to be highly representative of the *in vivo* bovine bronchial epithelium and can be utilised in the study of respiratory pathologies.

## Introduction

Through the process of inhalation, the airway epithelium is exposure to a wide variety of substances. These inhaled particles can potentially be harmful following contact, including pathogenic organisms and toxins [1–3]. As such, the respiratory tract is subject to diverse pathologies which can have numerous implications on both human and animal health. The airway epithelium is one of the first lines of defence, acting as a physiochemical barrier against inhaled toxins, pollutants and invading pathogens [2, 4]. The epithelia has developed numerous adaptions, including mucocilary clearance [5–7] and intercellular junctions [1, 8] which work in conjunction to remove inhaled insults, in order to maintain homeostasis [9]. These mechanisms can be disrupted following exposure by inhaled pathogens, which can cause extensive damage to the epithelia and allow transmigration to deeper tissue [10, 11]. The bronchial epithelium however is capable of repairing and remodelling itself following damage through proliferation and differentiation of progenitor cells, maintaining the integrity of the respiratory tract [12, 13]. Due to the impact of respiratory pathologies, there is a need to model the airway epithelium, including the associated defence mechanisms and the differentiation of epithelial cells during repair. This has traditionally been achieved through the use of animal models. Animal research is associated with high experimental cost and time requirements, as well as carrying ethical implications, due to the contradictions to the Three R’s principles [14]. As such, there is a need for physiologically-relevant *in vitro* models of the bronchial epithelium which can be used as alternatives to animal models.

Application of an air-liquid interface (ALI) to produce *in vitro* airway epithelial cell models (AECMs) is widely used in both toxicology [15–17] and infectious disease research [18–21]. These models trigger the differentiation of airway epithelial cells using exposure to the atmosphere [22], as well as through the addition of growth factors such as epidermal growth factor [23–25] and retinoic acid [23, 26, 27]. This results in a mixed-cell population consisting of ciliated cells, mucus-secreting goblet cells and progenitor basal cells present as a pseudostratified epithelium [28]. The differentiation process does not occur if primary cells are grown under submerged conditions [29], and as such submerged cultures fail to replicate the *in vivo* tissue complexity [30]. The transition of epithelial cells into a differentiated airway epithelium is complex process which occurs through a number of developmental stages, involving cell proliferation and differentiation [12, 31, 32]. Following differentiation, ALI-grown cultures possess both mucociliary clearance and transepithelial resistance [23, 32] which are critical for assessing their response to infection or toxic insult [33]. Differentiated AECM also enables infections to be studied in a mixed-cell population, allowing the identification of cell type targeting during infection [18, 31, 34, 35]. Due to these properties AECM provide excellent tools for researching respiratory pathologies.

The use of bovine airway epithelial cells to form differentiated AECM has been established previously. Bovine AECMs have been used to study economically-important infections in cattle, including bovine respiratory viruses [34, 36] and the bacteria *Mycobacterium tuberculosis* and *Mycobacterium bovis* [37]. Bovine AECMs have also been used to investigate the basic biology of the mammalian respiratory tract, including aspects of oxidative stress [38], ion transport and signalling [39, 40] and the air surface liquid [41]. The benefit of using cells isolated from cattle as opposed to human tissue is their ready availability and low cost [39]. As such, AECM derived from bovine airway epithelial cells may represent a more accessible alternative to human cells. This may be particularly relevant for infectious diseases in which identical or closely-related pathogens infect both humans and cattle. For example, *M. tuberculosis* is known to be carried by both humans and cattle [42]. Similarly, human respiratory syncytial virus is closely related to bovine respiratory syncytial virus, and results in similar associated pathologies [43].

For pathologies to be accurately assessed following exposure to infection or toxins, ALI models need to be fully characterised. Similarly, the transition of the model from undifferentiated cells to a fully differentiated epithelium must be well defined in order to determine the ability of the model to repair following damage. Although this has been achieved for human [28, 32] and ovine AECMs [44], bovine AECMs have yet to be well-defined. We aimed to fully characterise the differentiation of BBECs grown at an air-liquid interface. This model has previously been optimised in our group in order to establish a high degree of differentiation using a serum-free medium. Our bovine AECM was extensively studied at three day intervals, from three days prior to the establishment of an ALI, during which the cells were under submerged conditions, until day 42 post-ALI. Key markers of epithelial cell differentiation, including morphology, the formation of tight-junctions and the presence of ciliated and goblet cells were assessed at each time-point. De-differentiation and deterioration of the culture was also monitored, allowing us to define the optimum window at which the model was suitable used for infection or toxicology studies.

## Materials and Methods

### Isolation of bovine bronchial epithelial cells

Bronchial epithelial cells were obtained from cattle aged 18-36 months, as described in Cozens et al. The breeds of the animals used were Limousin (animal 1) and Simmental (animal 2 and 3). The lungs of cattle were collected post-slaughter at Sandyford Abattoir Ltd., UK. The bronchial tract was swabbed to ensure there was no pre-existing bacterial/fungal infection. Briefly, the main and lobar bronchi were isolated from the lungs and the surrounding tissue was carefully dissected away. Small (∼1 cm^2^) samples from each bronchus was collected and fixed in 2% (w/v) formaldehyde overnight for histological analysis and electron microscopy. The bronchi were sectioned and cut vertically to expose the epithelium, yielding rectangular tissue sections 6-7 cm in length. Epithelial cells were isolated from the bronchi by incubation overnight at 4°C in ‘digestion medium’ composed of Dulbecco’s modified Eagle’s medium (DMEM) and Ham’s nutrient F-12 (1:1) containing 1 mg/ml dithioreitol, 10 μg/ml DNAase and 1 mg/ml Protease XIV from *Streptomyces griseus*, supplemented with penicillin (100 U/ml), streptomycin (100 μg/ml) and amphotericin (2.5 μg/ml) (Sigma-Aldrich). All subsequent media used in this investigation were also supplemented with penicillin-streptomycin and amphotericin. The digestion was halted with the addition of foetal calf serum to give a final concentration of 10% (v/v). Loosely-attached epithelial cells were removed from the submucosa by rigorous rinsing of the luminal surface. The cell suspension was passed through a 70 μm cell strainer to remove tissue and was subsequently centrifuged and resuspended in ‘submerged growth medium’ (SGM), comprised of DMEM/Ham’s F-12 (1:1) supplemented with 10% (v/v) foetal calf serum. The viability of the cell suspension was assessed using the Trypan Blue exclusion assess, and was typically 90-95% viable. Cells were seeded into T75 tissue culture flasks (5 x 10^6^ cells/flask) for expansion. The flasks were incubated at 37°C in 5% CO_2_ and 14% O_2_, in a humidified atmosphere.

### Culture of bovine bronchial epithelial cells

The BBECs were harvested at 80-90% confluency (∼4 days). Cells were detached from the flasks using 0.25% trypsin-EDTA solution, centrifuged and resuspended in SGM at a density of 5 x 10^5^ cells/ml. The BBECs were seeded into the apical chamber of tissue culture inserts (Thincerts, Greiner #66540, polyethylene terephthalate membrane, 0.4 μm pore diameter, 1 x 10^8^ pore per cm^2^) at a cell density of 2.5 x 10^5^ cells per insert, and 1 ml of SGM was added to the basolateral compartment. Cultures were incubated at 37 °C, 5% CO_2_, 14% O_2_, in a humidified atmosphere. The BBECs were allowed to attach to the insert overnight. The apical medium of the culture was subsequently removed and the apical surface washed with 0.5 ml PBS. The SGM media in the apical and basolateral compartments was then replaced. This process was repeated every 2 – 3 days. The trans-epithelial electrical resistance (TEER) of the cultures were monitored on a daily basis using an EVOM2 epithelial voltohmmeter (World Precision Instruments, UK), as per the manufacturer’s instruction. The SGM was replaced with a mixture of SGM and ‘air-liquid interface medium’ (ALIM) (1:1) when the TEER reached above 200 Ω/cm^2^ (∼2 days post-seeding). The ALIM was composed of DMEM and airway epithelial cell growth medium (Promocell) (1:1) supplemented with 10 ng/ml epidermal growth factor, 100 nM retinoic acid, 6.7 ng/ml triiodothyronine, 5 μg/ml insulin, 4 μl/ml bovine pituitary extract, 0.5 μg/ml hydrocortisone, 0.5 μg/ml epinephrine and 10 μg/ml transferrin (all Promocell). When the TEER value was above 500 Ω cm^2^ (∼6 days post-seeding), an ALI was generated by removing the medium in the apical compartment, thereby exposing the epithelial cells to the atmosphere (day 0 post-ALI). Following the formation of the ALI, the cells were fed exclusively from the basal compartment with ALIM. Apical washing, basal feeding and TEER measurements were performed every 2 - 3 days until day 42 post-ALI.

### Histology

Samples of BBEC cultures were taken at three day intervals, from three days prior to the establishment of an ALI (day −3) until day 42 post-ALI. At each time-point, samples were fixed by the addition of 4% (w/v) paraformaldehyde to the apical surface for 15 min at room temperature. Fixed samples were rinsed and stored in PBS at 4°C until the completion of the time-course. A series of increasing ethanol concentrations was used to dehydrate the samples, before being cleared with xylene, infiltrated with paraffin wax and embedded in wax blocks. Sections of 2.5 μm thickness were cut in transverse sections using a Thermoshandon Finesse ME+ microtome. Samples were subsequently haematoxylin and eosin (H&E) stained using standard histological techniques.

### Histological analysis

Histological sections stained with H&E were prepared from three individual cultures derived from each of three animals. For each section, five randomised x400 fields of view were imaged across the stand. ImageJ was used to quantify the thickness of the cell layer and the number of cell layers forming the epithelium at three vertical sections within each field of view. In addition, the number of ciliated cells (cells with visible cilia present), epithelial gaps and pyknotic cells were quantified within each field of view.

### Chromogenic Immunohistochemistry

For the identification of basal cells, chromogenic immunohistochemistry was used. Samples were processed and 2.5 μm-thick sections obtained as described. A Menarini Access Retrieval Unit was used to perform heat-induced epitope retrieval. Staining was subsequently performed using a Dako Autostainer. Endogenous peroxidase was blocked using 0.3% (v/v) H_2_O_2_ in PBS. Basal cells were identified by incubation for 30 min with a 1:30 dilution of mouse anti-p63 antibody (Abcam; #ab735), application of an anti-mouse HRP-labelled polymer and visualization with a REAL EnVision Peroxidase/DAB+ Detection System (Dako; #K3468). Samples were counterstained with Gill’s haematoxylin, before dehydration, clearing and mounting in synthetic resin. Sections were viewed using a Leica DM2000 microscope (Leica, Germany).

### Fluorescent Immunohistochemistry

Samples were processed and 2.5 μm-thick sections obtained as described. Samples were deparaffinised using two 5 min washes in 100% xylene before rehydration through a series of decreasing ethanol concentrations. Samples were subject to heat-induced epitope retrieval by immersion in sodium citrate buffer (10 mM sodium citrate, 0.05% [v/v] Tween-20, pH 6) at 100 °C for 20 min. Samples were blocked by incubation in blocking buffer for 1 h at room temperature (PBS containing 0.05% [v/v] Tween-20, 10% [v/v] goat serum and 1% [w/v] bovine serum albumin). The cultures were incubated with primary antibodies diluted in blocking buffer for 1 h at room temperature. Samples were washed three times in PBS containing 0.05% (v/v) Tween-20 for 2 min following each incubation. The primary-secondary antibody pairings were applied as follows. Primary antibodies were used at a dilution of 1:200. Secondary antibodies were used at a dilution of 1:400. Ciliated cells were detected with rabbit anti-β-tubulin antibody (Abcam; #ab6046) and visualised using goat anti-rabbit-Alexa Fluor 647 (Thermo Fisher; #A-21244); basal cells were visualised with mouse anti-p63 antibody (Abcam, #ab735) and visualised with goat anti-mouse-Alexa Fluor 568 (Thermo Fisher; #A-11031); goblet cells were detected with fluorescein-labelled jacalin (Vector Laboratories; FL-1151) [45]. Following each primary-secondary pairing blocking was repeated. Nuclei were stained with 300 nM 4’,6 diamidino-2-phenylindole (DAPI) for 10 min. Samples were subsequently washed and mounted in Vectashield mounting medium (Vector Laboratories). Samples were observed on a Leica DMi8 microscope. Analysis of captured images was performed using ImageJ software.

### Immunofluorescence microscopy

For immunofluorescence, paraformaldehyde-fixed samples were permeablised using permabilization buffer (PBS with 0.5% [v/v] Triton X-100, 100 ml/ml sucrose, 4.8 mg/ml HEPES, 2.9 mg/ml NaCl and 600 μg/ml MgCl_2_, pH 7.2) for 10 min at room temperature. Samples were blocked by incubation with blocking buffer for 1 h. Ciliated- and goblet-cells were detected using the methods described for fluorescent immunohistochemistry. Tight-junction formation was detected with mouse anti-ZO-1 antibody (1:50 dilution; Thermo Fisher; #33910) and visualised with goat anti-mouse-Alexa Fluor 488 (1:400 dilution; Thermo Fisher; #A-11001). Antibodies utilised were diluted in blocking buffer. The cultures were incubated with primary antibodies diluted in blocking buffer for 1 h at room temperature. Samples were washed three times in PBS containing 0.05% (v/v) Tween-20 for 2 min following each incubation. F-actin was visualised by incubation with a 1:40 dilution of rhodamine phalloidin (Thermo Fisher; #R415) for 20 min. Nuclei were stained with 300 nM DAPI for 10 min. Following staining, membranes were cut from their insert and mounted in Vectashield mounting medium (Vector Laboratories). Samples were observed on a Leica DMi8 microscope. Analysis of captured images was performed using ImageJ software.

### Quantification of ciliogenesis

To quantify the degree of cilia formation on the apical surface, five randomized fields of view of each β-tubulin-stained insert were acquired via a 20x objective. Coverage of cilia was quantified for each image using ImageJ. A fluorescence intensity threshold was applied to select ciliated regions. These regions were measured and expressed as a percentage of the total area.

### Scanning electron microscopy

Cultures were fixed in 1.5% (v/v) glutaraldehyde diluted in 0.1 M sodium cacodylate buffer for 1 h at 4 °C. The apical and basal compartment was subsequently rinsed three times and stored in 0.1 M sodium cacodylate buffer at 4 °C until completion of the time-course. To post-fix samples, 0.5 ml of 1% (w/v) osmium tetroxide was added to the apical compartment for 1 h at room temperature and washed three times for 10 min with distilled water. Samples were stained with 0.5% (w/v) uranyl acetate for 1 h in the dark. Dehydration was performed using a series of increasing ethanol concentrations. Hexamethyldisilazane was used for the final drying stage before being placed in a desiccator overnight. Membranes were cut from the inserts, mounted onto aluminium SEM stubs and gold sputter-coated. Samples were analysed on a Jeol 6400 scanning electron microscope at 10 kV.

### Transmission electron microscopy

For transmission electron microscopy (TEM), samples were fixed and processed as described for scanning electron microscopy until dehydration in absolute ethanol. Samples were subsequently washed in propylene oxide three times for 10 min before being immersed in 1:1 dilution of propylene oxide and Aridite/ Epon 812 resin overnight. The samples were washed in three changes of resin and fresh embedded in resin within rubber models, which was allowed to polymerise at 60 °C for 48 h. Resin-embedded samples were cut to ultrathin sections of 50 nm thickness using a Leica Ultracut UCT and a DiATOME diamond knife. Sections were collected on 100 mesh Formvar-coated copper grids. Samples were finally contrast stained with 2% (w/v) methanolic uranyl acetate for 5 min and Reynolds lead citrate for 5 min. Cultures were analysed on a FEI Tecnai transmission electron microscope at 200 kV. Images were captured with a Gatan Multiscan 794 camera.

## Results

### Epithelial morphology

Bovine bronchial epithelial cells were cultured at an ALI over 42 days. Morphological assessment of the epithelium was conducted on histological sections taken from samples fixed at three day intervals, ranging from three days prior to establishment of the ALI (day −3 pre-ALI) until day 42 post-ALI. The general morphology of the epithelial cell layer was assessed using an H&E stain (Fig 1A; S1 Fig). During submerged growth (day −3 and 0), BBECs formed squamous monolayers which exhibited no evidence of polarisation (Fig 1A [i]). Establishing an ALI caused the cultures to transition to a differentiated pseudostratified epithelium over time, reminiscent of *ex vivo* tissue (S1 Fig). Between day 0-21 post-ALI the cell layer gradually thickened to approximately 30-40 μm thickness. Conversely, there was no subsequent increase in the thickness of the cultures from day 21 onwards (Fig 1C). The number of cells within the epithelium also increased following the establishment of the ALI (Fig 1D). The BBECs transitioned from a squamous monolayer to becoming approximately two cells in depth and possessing a cuboidal morphology between day 3-12 post-ALI (Fig 1A [ii]). By day 15, until completion of the time-course, the cultures were approximately three cells in depth. This change coincided with the epithelium becoming increasingly columnar and pseudostratified (Fig 1A [iii & iv]), replicating *ex vivo* tissue (Fig 2A [i]). At all time-points from day 12 to day 42, a single layer of p63 positive basal cells was observed. This mimicked the distribution of basal cells in *ex vivo* samples where a single continuous layer was present attached to the basement membrane (Fig 1B; S2 Fig).

**Figure 1.**
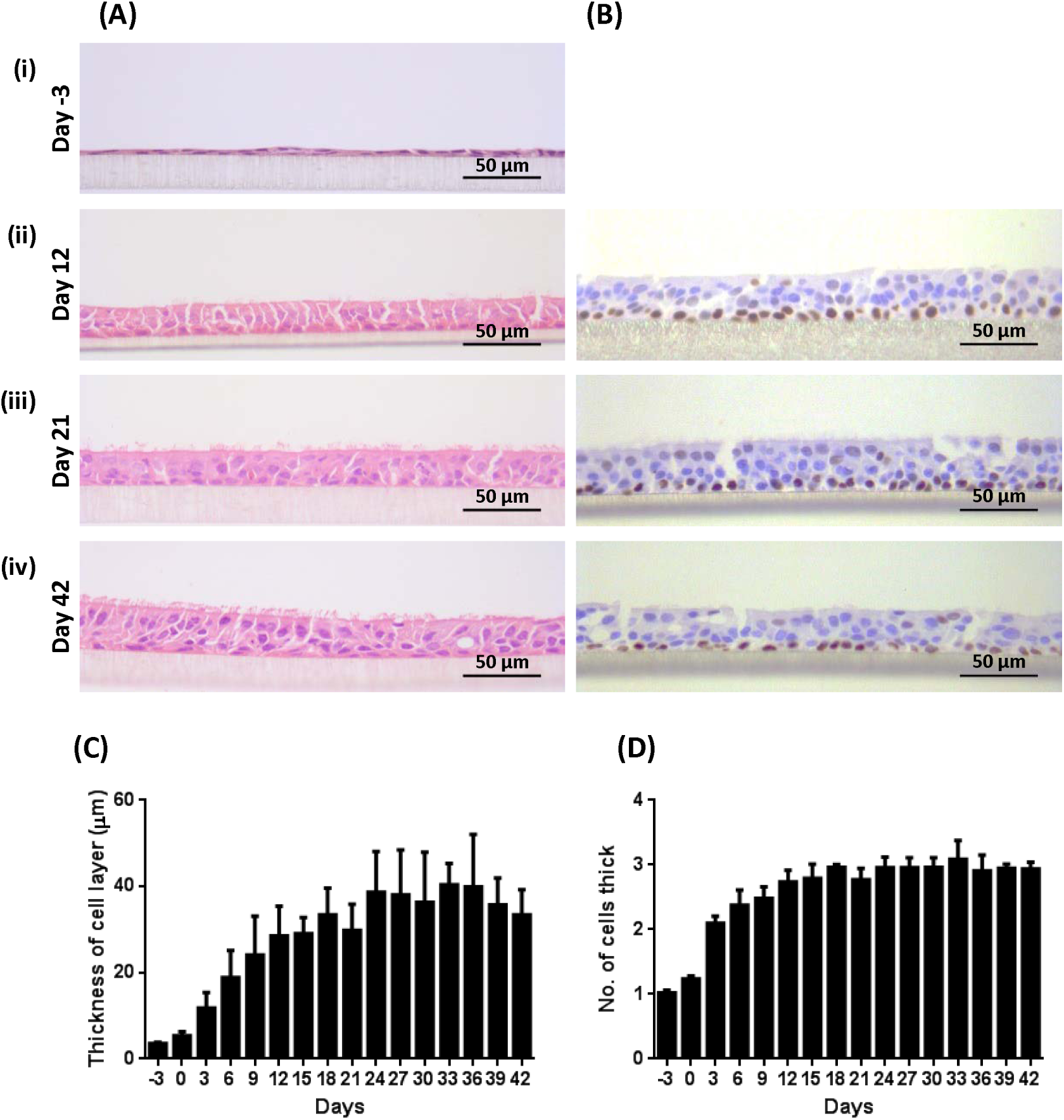
Histological assessment of the differentiation of BBEC cultures over time. BBEC cultures were grown for a stated number of days at an ALI before being fixed, paraffin-embedded and sections cut using standard histological techniques. Sections were subsequently deparaffinised and stained using (A) H&E stain and (B) immunohistochemistry with an anti-p63 antibody labelling the nuclei of basal cells. Representative images are shown of (i) day −3, (ii) day 12, (iii) day 21 and (iv) day 42 post-ALI (see S1 Fig). Quantitative analysis (using ImageJ) of histological sections of BBEC layers fixed at three day intervals ranging from day −3 pre-ALI to day 42 post-ALI (see S1 Fig), showing (C) epithelial thickness and (D) the number of cell layers composing the epithelium was performed. For each insert, three measurements were taken (left, centre and right) in each of five 400x fields of view distributed evenly across the sample; three inserts were analysed per time point and the data represents the mean +/-standard deviation from tissue derived from three different animals.

The BBEC cultures were assessed for cellular and tissue deterioration following extended culturing at an ALI. There was a time-dependent increase in both the number of pyknotic cells (S3A [i] & B Fig) and epithelial gaps (S3A [ii] & C Fig) present per field of view. However this trend was determined to be due to an overall increase in the number of cells present in the epithelium over time. Once the cell number in the epithelium had reached a peak at day 21 post-ALI, there was no subsequent significant increase in either the number of epithelial gaps or pyknotic cells (Ordinary one-way ANOVA). This finding suggested the BBECs were stable for at least six weeks of culturing at an ALI.

A comparison was made of histological sections taken from BBECs cultured for 21 days post-ALI and the *ex vivo* bovine bronchial epithelium of the donor animal (Fig 2). Both sections display a pseudostratified columnar epithelial morphology, stereotypical of the airway epithelium (Fig 2A). The BBECs grown at an ALI were consistently thinner in comparison to the *ex vivo* epithelium. Immunohistochemistry was performed to detect the distribution of the epithelial cell types of the lower respiratory tract (Fig 2B). Both *ex vivo* tissue and the BBECs grown at an ALI displayed all major epithelial cell types with comparable morphology. B-tubulin-labelled ciliated cells and jacalin-labelled mucus producing goblet cells were present at the apical aspect of the epithelium; however the density of ciliated cells was lower in the cultured BBECs compared to *ex vivo* samples. The differentiation of ciliated and mucus producing epithelial cells will be discussed in greater detail below.

**Figure 2.**
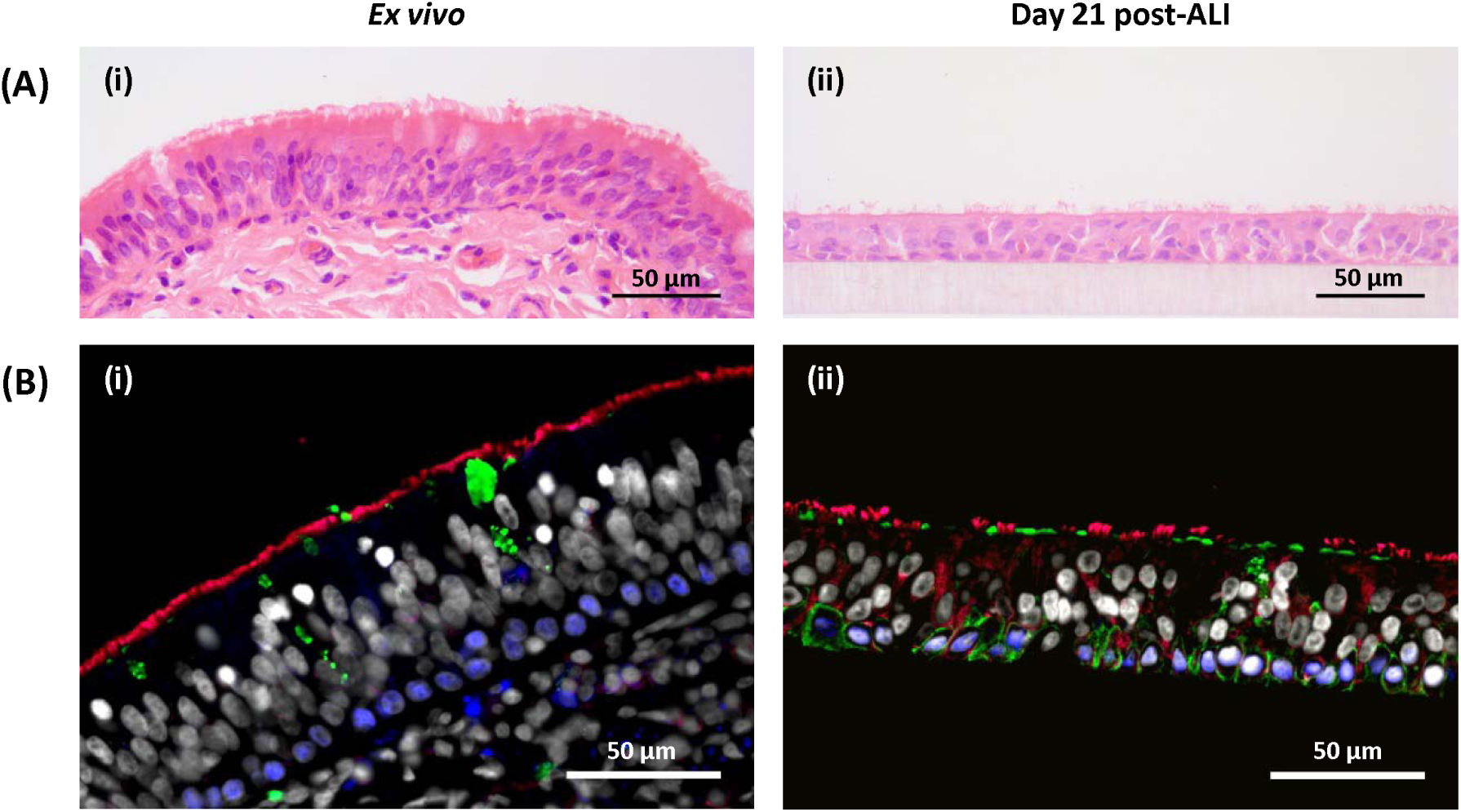
Comparison of epithelial morphology and distribution of cell types between differentiated BBECs and the bovine bronchial epithelium. BBEC cultures were grown for 21 days at an ALI before being fixed, paraffin-embedded and sections cut using standard histological techniques; sample of *ex vivo* tissue were also taken from the donor animal. Sections were subsequently deparaffinised and stained using (A) H&E stain and (B) immunohistochemical stain for epithelial cell type markers (β-tubulin - red; Muc5AC - green; p63 - blue; nuclei - grey). Representative images are shown of (i) *ex vivo* bovine bronchial epithelium and (ii) differentiated BBECs 21 days post-ALI.

### Barrier function

The barrier function of the BBECs was assessed at three day intervals during culturing (Fig 3). Using junctional protein ZO-1 as a marker, tight-junctions were found to be present in the BBEC cultures at all time-points, both during submerged and ALI growth (Fig 3A and S3). ZO-1 was shown to be localised to the sub-apical cell-to-cell borders, indicative of intact tight-junction. During submerged growth (day −3 and 0) the cells were large and squamous, but once an ALI was established the cells adopted a more cobblestone appearance, reminiscent of differentiated epithelia, and as such the number of tight-junctions present per field of view increases (Fig 3A). There was no detectable decrease in ZO-1 at later time-points. Transmission electron microscopy of day 42 post-ALI cultures and *ex vivo* further identified adherens junctions and desmosomes (Fig 3B), confirming the presence of junctional complexes.

**Figure 3.**
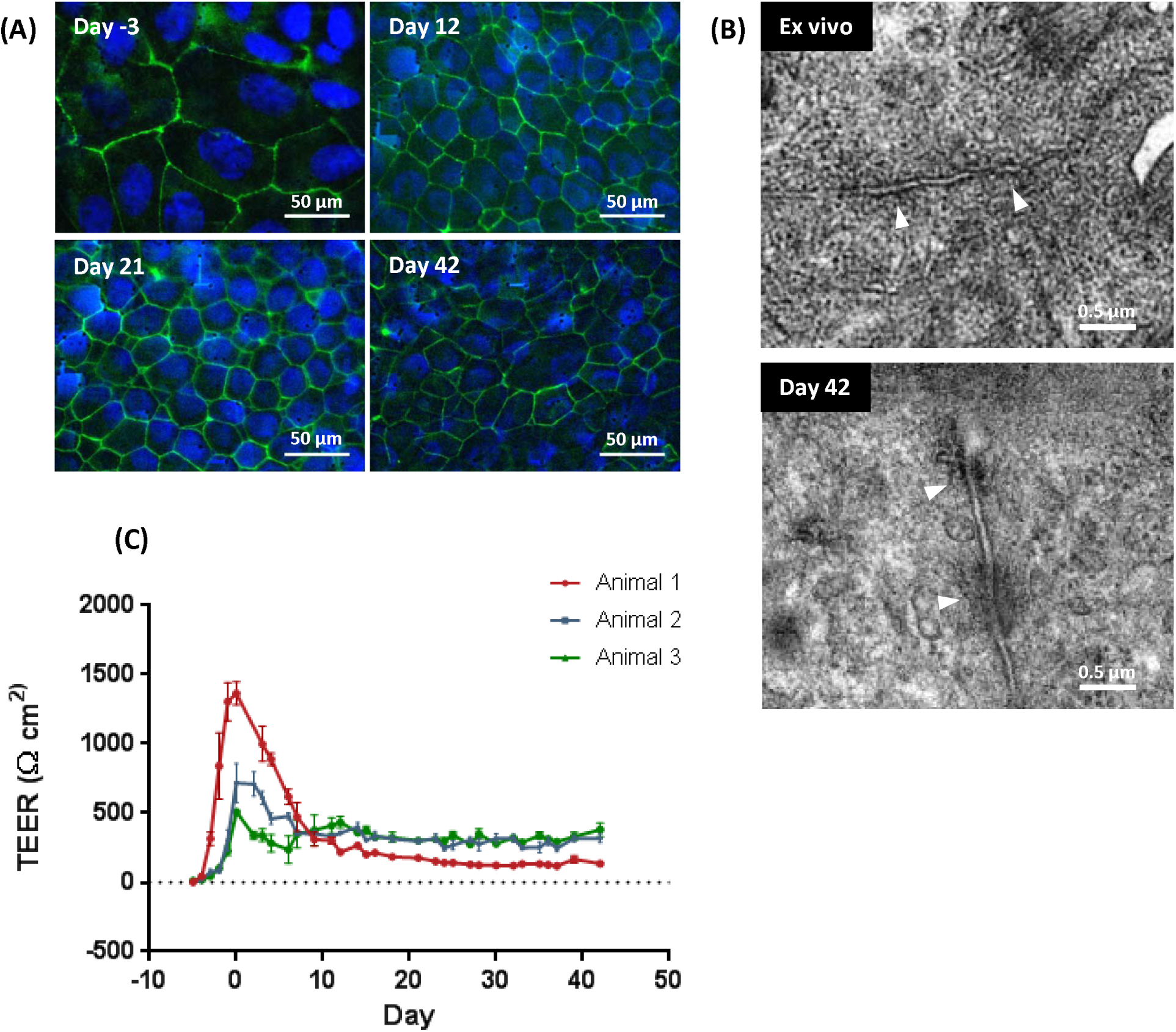
Barrier function and tight-junction formation in BBEC cultures over time. BBEC cultures were grown for the stated number of days at an ALI before fixation. Tight-junction formation of the BBEC cultures was subsequently assessed using (A) immunofluorescence imaging of tight-junctions (ZO-1 - green; nuclei - blue). Representative images are shown of (i) day −3, (ii) day 12 (iii), day 21 and (iv) day 42 post-ALI (see S4 Fig). Transmission electron micrographs in (B) display the presence of junctional complexes (arrowheads denote adherens junctions and desmosomes). Representative images are shown of (i) *ex vivo* bovine bronchial epithelium and (ii) differentiated BBECs 42 days post-ALI. Tight-junction integrity during the course of culturing was assessed by measuring the TEER of BBEC cultures. Nine inserts were analysed per growth condition and the data represents the mean +/-standard deviation from tissue derived from three different animals.

The TEER of the BBECs was measured to confirm the function of tight-junctions in the epithelium (Fig 3C). The BBECs reached confluency after two days of submerged growth post-seeding, resulting in the presence of a TEER. The TEER peaked after five days of submerged growth in all replicates; however the value at which the TEER peaked varied considerably between replicates. At this time-point, barrier function was present and an ALI could be established. The TEER gradually declined thereafter during the ALI phase, until day 9 post-ALI, by which TEER stabilised at ∼150-300 Ω × cm^2^ in all replicates. The reduction in TEER did not coincide with variation in tight-junction staining (Fig 3A), and barrier function was intact throughout the 42 days of culturing at an ALI.

### Cilia formation

The temporal development of cilia on the apical surface of the AECM was assessed during culturing (Fig 4). Cilia were identified using β-tubulin as a marker (Fig 4A; S4 Fig), detection using SEM (Fig 4B; S5 Fig) and in histological sections (Fig 4C; S1 Fig). Furthermore, the extent of cilia formation was quantified in histological samples (Fig 4D) and immunostained cultures (Fig 4E). During submerged growth, β-tubulin could be detected in BBECs, however the staining pattern was cytoplasmic, forming cytoskeletal microtubules, and was not indicative of cilia formation (Fig 4A [i]). This was confirmed by SEM (Fig 4B [i]). Cilia formation was evident as early as day 6 post-ALI; cilial staining was distinguished from cytoskeletal β-tubulin due to the intensity of the signal and localisation at the apical aspect (Fig S5). As cells were cultured over time, cilia formation at the apical aspect became increasingly abundant. Bright-field microscopy of BBECs grown at an ALI for 21 days on low-pore density inserts showed that cilia are capable of beating microspheres in a coordinated fashion (data not shown), confirming the cilia are functional. The degree of cilia formation of the BBECs reached a maximum by approximately day 21 post-ALI, and there is no subsequent significant increase in the number of ciliated cells between day 21-42 post-ALI (Figs 4D and 4E; Ordinary one-way ANOVA). During this period the majority of the apical aspect was composed of ciliated cells (Fig 4A [iii & iv]). The time by which maximal cilia formation was achieved varied between cultures derived from individual animals, however the overall temporal pattern of cilial differentiation was similar (Fig 4D and 4E), and variation was not statistically significant (Ordinary one-way ANOVA).

**Figure 4.**
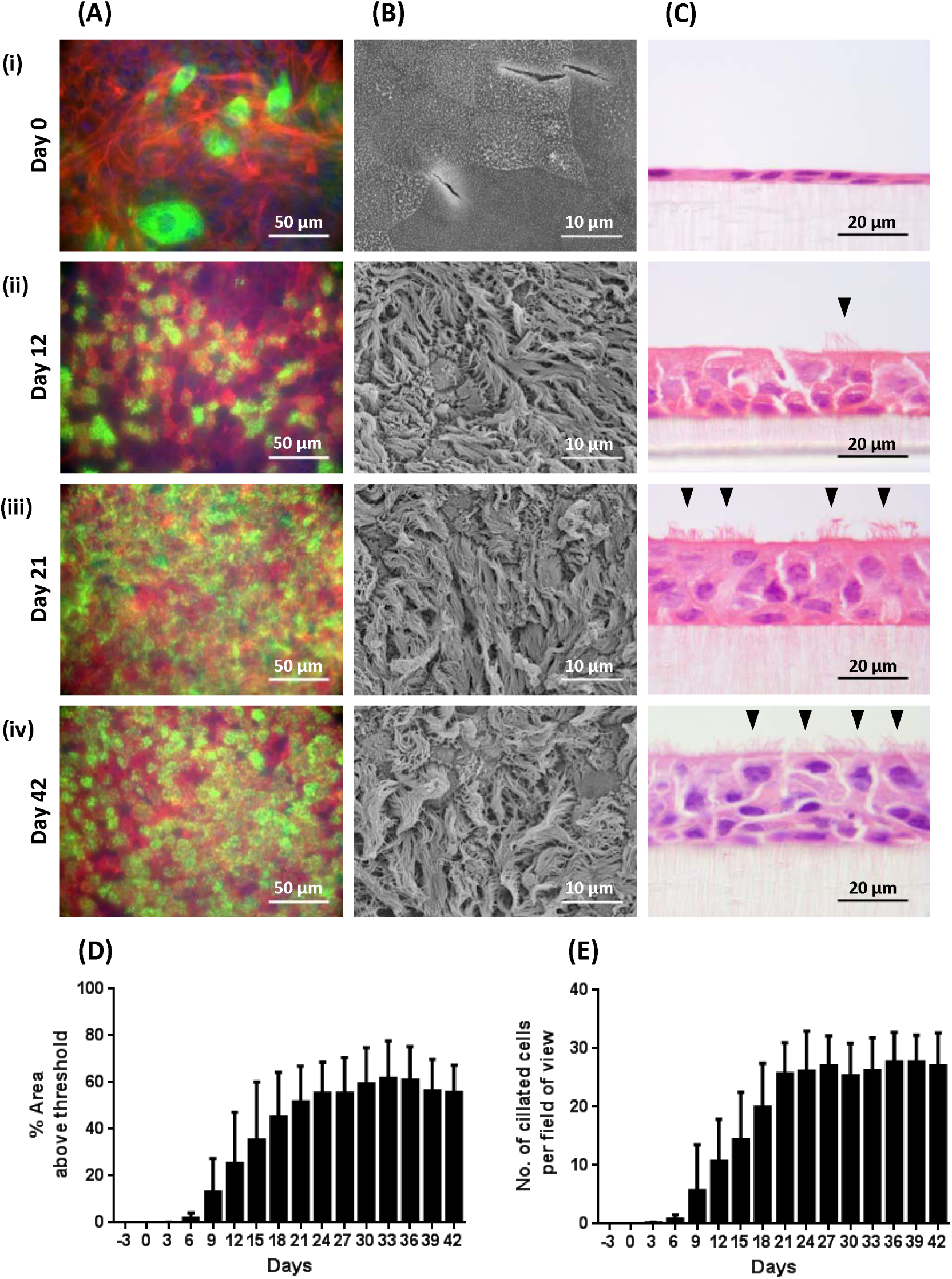
Cilia formation in BBEC cultures over time. BBEC cultures were grown for the stated number of days at an ALI before fixation. The BBEC cultures were subsequently processed to assess cilia formation using (A) immunofluorescence labelling of cilia (β-tubulin - green; F-actin - red; nuclei - blue), (B) examination by SEM (arrowheads denote ciliated cells) and in (C) H&E stained histological sections. Representative images are shown of (i) day 0, (ii) day 12, (iii) day 21 and (iv) day 42 post-ALI (see S1, S5 & S6 Fig). Quantitative analysis of cilia formation (using ImageJ) of BBEC cultures fixed at three day intervals ranging from day −3 pre-ALI to day 42 post-ALI using (D) fluorescence intensity thresholding of immunostained cultures (see S5 Fig) and (E) by counting the number of ciliated cells per field of view in H&E-stained sections (see S1 Fig). In (D), cilia formation was quantified by measuring the area above a fluorescence intensity threshold in ImageJ; for each insert, five regions evenly distributed across the sample were measured. In (E), for each insert, ciliated cells were counted in each of five 400x fields of view evenly distributed across the sample. For all of the above quantifications, three inserts were analysed per time point and the data represents the mean +/- standard deviation from tissue derived from three different animals.

Cilia formation was compared between day 21 post-ALI BBEC cultures and *ex vivo* tissue (Fig 5). Both samples displayed a highly ciliated apical surface. However the overall degree of coverage was slightly lower in BBEC cultures grown at an ALI in comparison to *ex vivo* tissue (Fig 5A). The ultrastructure of the cilia produced by the BBEC cultures was highly analogous to the source tissue (Fig 5B and 5C). The configuration of both the cilium basal body (Fig 5B) and the ciliary 9 + 2 axoneme (Fig 5C) was consistent; there was no indication of malformation in cilia produced by BBEC.

**Figure 5.**
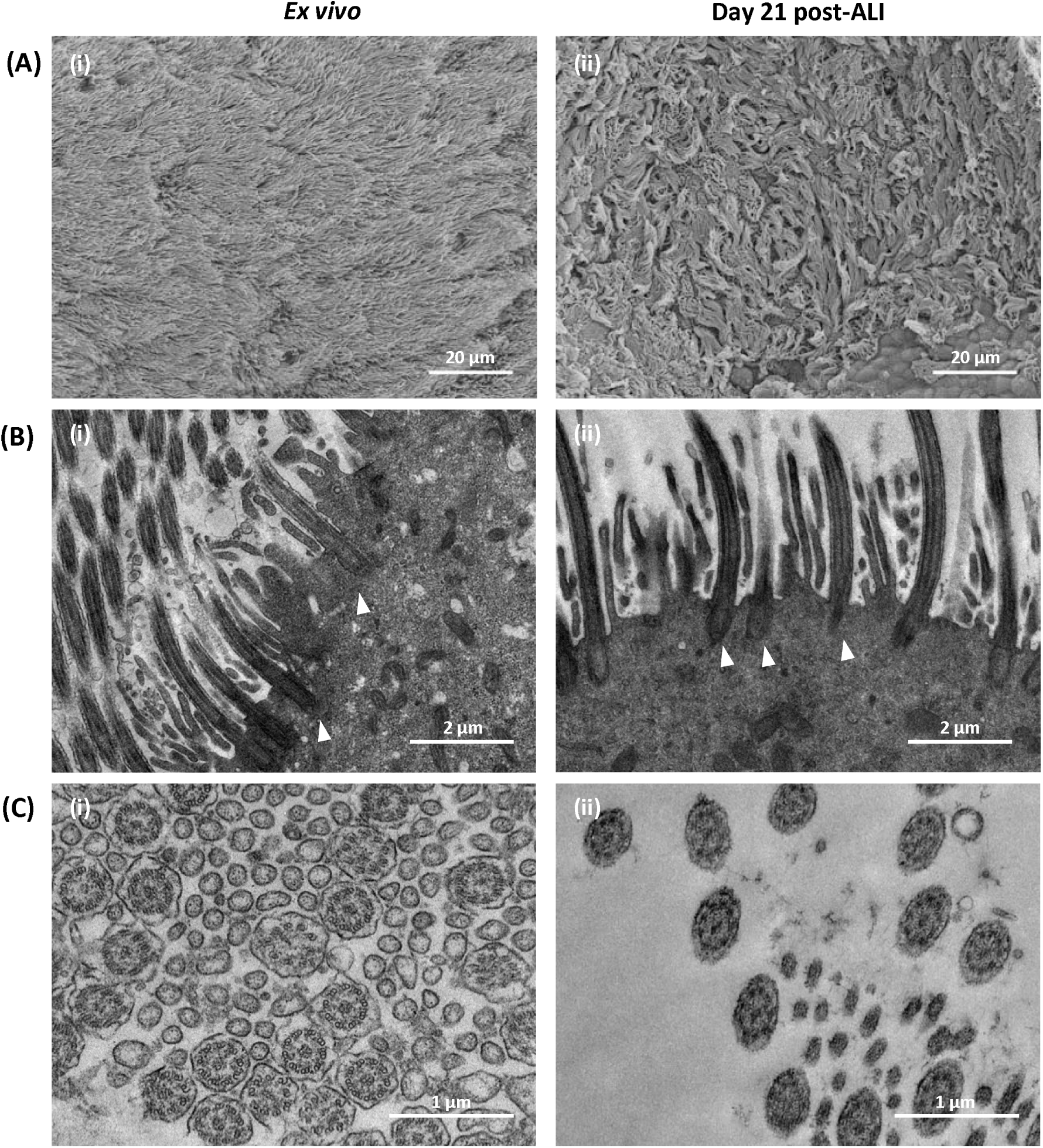
Electron microscopy of cilia formation in differentiated BBECs compared with bovine bronchial epithelium. BBEC cultures were grown for 21 days at an ALI befor being fixed and processed for electron microscopy, a sample of *ex vivo* tissue were also taken from each donor animal. Images are shown of (A) scanning electron micrographs of apical surface, (B) transmission electron micrographs of cilium basal bodies (arrowheads denote basal bodies) and (C) transmission electron micrographs of 9 + 2 axoneme of cilia. Representative images are shown of (i) *ex vivo* bovine bronchial epithelium and (ii) differentiated BBECs 21 days post-ALI.

### Mucus Production

The development of mucus-producing goblet cells was also assessed in ALI-grown BBECs (Fig 6). Mucus production was identified using jacalin, a lectin which binds mucin Muc5AC on goblet cells [28] (Fig 6A; S7 Fig). Muc5AC-positive cells were present in BBEC cultures, both at an ALI and whilst cells were submerged. There was no observable change in the number of Muc5AC-positive cells over time. Scanning electron microscopy further confirmed the presence of mucus in the BBEC models. Excreted mucus could be observed in cultures from day 15 post-ALI, either as globules coating cilia within the model (Fig 6C [i]), as well as webs coating the apical surface (Fig 6C [ii]). Goblet cells actively extruding mucus could also be observed (Fig 6C [iii]). This suggests the mucosal phenotype of the BBECs was not dependent on full differentiation of the model, however active extrusion of mucus could only be confirmed from day 15 post-ALI onwards.

**Figure 6.**
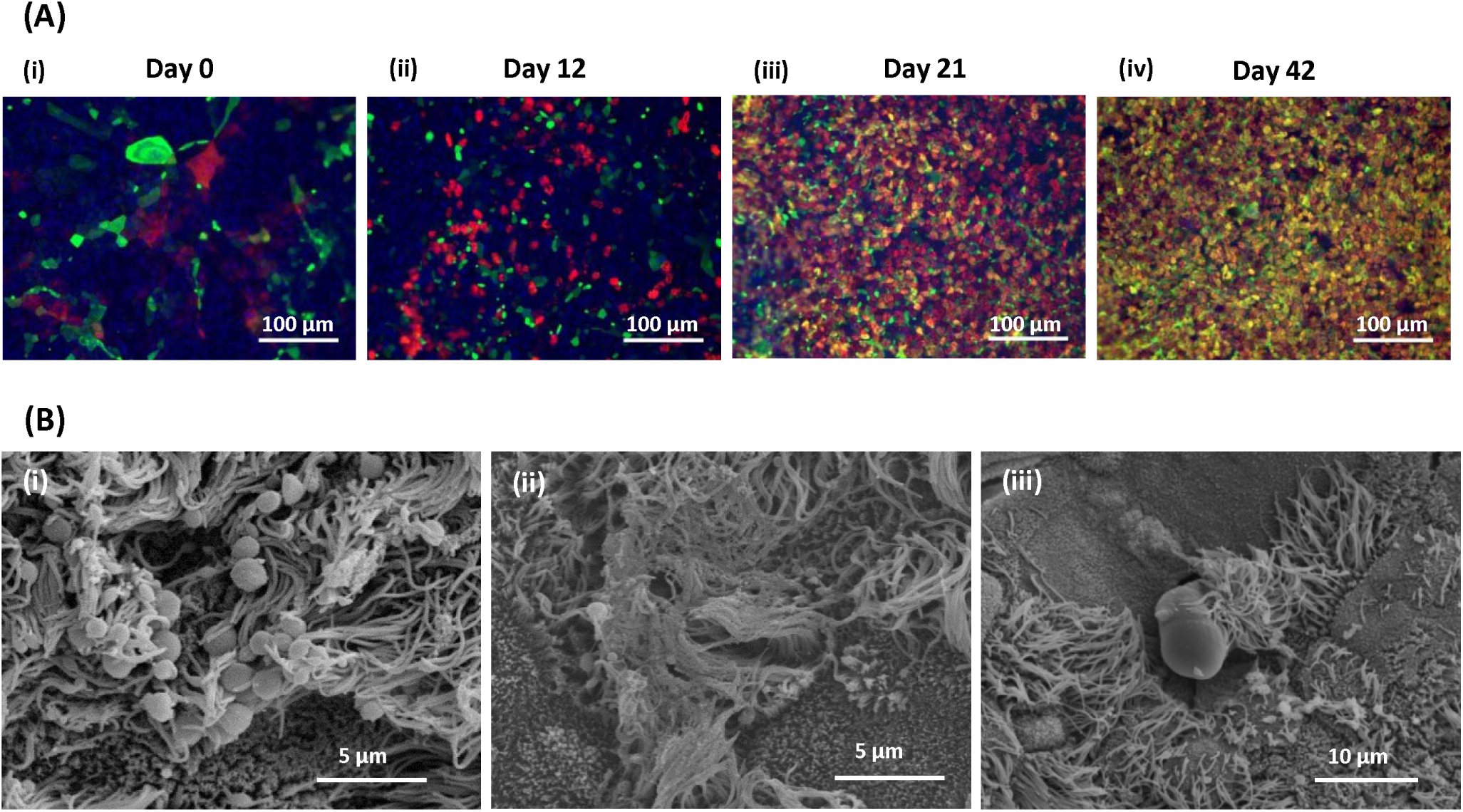
Mucus production in BBEC cultures over time. BBEC cultures were grown for the stated number of days at an ALI before fixation. The BBEC cultures were subsequently processed to assess mucus production using (A) immunofluorescence imaging of mucus formation (Muc5AC - green; β-tubulin - red; nuclei - blue). Representative images are shown of (i) day 0, (ii) day 12, (iii) day 21 and (iv) day 42 post-ALI (see S8 Fig). The presence of mucus on the apical surface was also imaged in (C) scanning electron micrographs of BBEC cultures grown at an ALI. Representative images are shown of (i) globules of mucus coating cilia (day 33 post-ALI), (ii) web of mucus coating the apical surface (day 36 post-ALI) and (iii) mucus extruded by a goblet cell (day 21 post-ALI).

### Ultrastructure

Analysis of the ultrastructure of the bovine AECM was conducted at each three day interval using SEM. During submerged growth, the cells appeared squamous (Fig 7A) and devoid of markers of differentiation such as cilia. Microvilli and microplicae could be observed on the cellular surface (Fig 7B). Once an ALI was introduced, cells appear to form a more cobblestone morphology. This was accompanied by the microvilli becoming denser and more pronounced (Fig 7C). Cilia were visible by day 6 post-ALI in isolated cells (Fig 7D). Ciliated cells became more numerous as the model progresses through the differentiation period (day 6-21 post-ALI) (Fig 7E). There was no observable difference in the topography of the model between days 21-42 post ALI, with little sign of degradation or dedifferentiation in the cell culture present visually after 6 weeks of culturing (Fig 7F). Cross sections of the cell layer at day 18 post-ALI exhibited the stereotypical pseudostratified morphology (Fig 7G). In differentiated BBEC cultures, the majority of cells at the apical aspect were ciliated (Fig 5A [i]), with microvilli observed around the base of the cilia (Fig 7I). Highly microvillous cells were also present, which may be non-excreting goblet cells (Fig 7H) [46]. Globules of mucus located nearby and extruding mucus could be observed in the model, supporting this assumption (Fig 6C).

**Figure 7.**
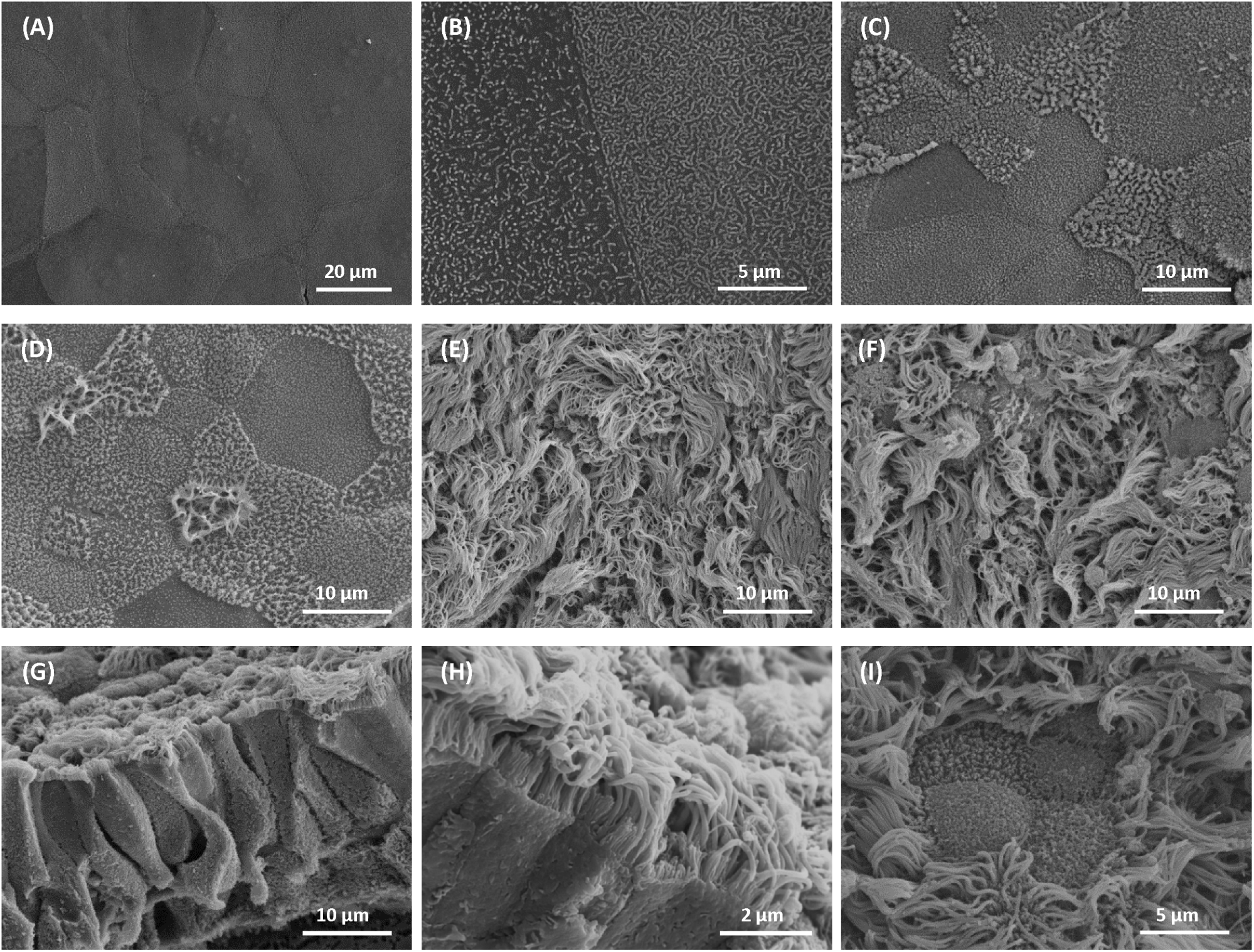
Electron microscopic assessment of the ultrastructure of BBEC cultures over time. BBEC cultures were grown for the stated number of days at an ALI before fixation and processing for SEM. Scanning electron micrographs of cultures are shown of representative images of (A) the undifferentiated apical surface (day 0 post-ALI), (B) microvilli and microplicae on undifferentiated cells (day 0 post-ALI), (C) microvilli on differentiating cell (day 6 post-ALI), (D) an early ciliated cell (day 6 post-ALI), (E) the differentiated apical surface (day 18 post-ALI), (F) the differentiated apical surface (day 42 post-ALI), (G) the pseudostratified epithelium (day 18 post-ALI), (H) microvilli on ciliated cells (day 36 post-ALI) and (I) non-extruding goblet cells (day 21 post-ALI).

### Discussion

The aim of the present study was to characterise the differentiation over time of an AECM derived from BBECs. The model was assessed over a 42-day period to allow the identification a window at which the culture was at an optimum differentiation. This has important implications for the use of the model in respiratory infection experiments. The degree of differentiation of airway epithelial cells can considerably alter the ability of both bacterial and viral pathogens to colonise [25, 31, 47]. Differentiation state of an AECM can also impact the response of a model during both infectious and toxicology studies [48, 49]. It was as such vital to pinpoint the window at which our bovine AECM is fully differentiated. Previous studies have placed this window between day 24-33 for human AECMs [32] and day 24-42 for ovine AECMs [44]. We have carried out comprehensive analysis of the differentiation of our model over an extended time-period and determine the window in which the bovine AECM were fully differentiated as between day 21-42 post-ALI.

Within the bovine AECM, the major cell types present in the bronchial epithelium (ciliated-, goblet- and basal cells) were replicated in a pseudostratified morphology comparable to *ex vivo* tissue (Fig 2B). The epithelium did not exhibit evidence of dedifferentiation of these cell-types over the six weeks of culture at an ALI; there was no reduction in either the number of ciliated- or goblet cells by day 42 post-ALI (Figs 4 & 6). The model was further assessed for signs of degradation. There was no significant increase in the number of pyknotic cells or epithelial gaps between day 15 (at which the cell morphology had reached peak thickness) and day 42 post-ALI (S3 Fig; Ordinary one-way ANOVA). This suggests there was no observable increase in cell death in the AECM, either due to apoptosis or autophagy [50, 51] following extended periods of culturing. Importantly, this confirms that once fully-differentiated, the model was stable for extended periods. A single culture can therefore be utilised in experimentation for up to 21 days, making the bovine AECM suitable for long-term or concurrent infections or repeat exposure toxicology studies.

The formation of tight-junctions between cells creates a physiochemical barrier against inhaled substances and prevents the penetration of pathogens or chemicals into the interstitial compartment [1, 8, 52]. Infection with certain viruses or bacteria however can cause transient disruption of tight and adherens junctions [11, 47, 53], and as such their presence is an essential feature of AECM for modelling pathologies. The TEER is an important method for assessing the formation of junctional complexes between cells [54]. In our model, once confluency was reached, the TEER of the culture rapidly increased (Fig 3C), suggestive of the formation of junctional complexes. The TEER peaked after approximately five days of submerged culturing, and this coincided with the formation of barrier function within our model. There was variation between animals in the peek value of TEER. Once an ALI was established, TEER decreased and stabilised at ∼150-300 Ω × cm^2^ by day 9 post-ALI. There was no subsequent decrease in TEER up to day 42 post-ALI. This trend in TEER replicated the pattern exhibited in cultures derived from other animals [25, 44, 55], however TEER of other AECMs have been shown to stabilised at the peak [32]. This variation in TEER over the course of culturing was not reflected in the presence of tight-junctional protein ZO-1, which was maintained at a stable level throughout the time-course. ZO-1 localised towards the cell-cell borders at all time-points (Fig 3A; S4 Fig), suggesting the presence of tight-junctions in the model both during submerged and ALI growth. As expected, the tight-junctions localised at the subapical region [8]. Using TEM, the structure of both desmosomes and adherens junctions of the bovine AECM were also observed, and was highly representative of *ex vivo* junctional complexes, even at the end point of the time-course (Fig 3B). This evidence suggests our model possesses stable junctional complexes, allowing the effect of disruption of these barriers following challenge from either pathogens or toxin to be defined.

A highly ciliated apical surface is one of the hallmarks of the respiratory epithelium. Ciliated cells work in conjunction with mucus production to ensure removal of invading pathogens and inhaled particles through the process known as mucociliary clearance [56]. This occurs through a system which entraps inhaled objects in globules of mucus which are subsequently swept from the respiratory epithelium by the coordinated movement of cilia [6]. This mechanism is the first line of defence of the respiratory epithelium, and as such its presence is a vital component of AECMs [57]. Cilia formation was detected and quantified in BBECS using immunofluorescence microscopy and in histological sections (Fig 4D and E). Cilia were present in our model from as early as day 6 post-ALI, sooner than previous models have reported [31, 32]. Ciliated cells significantly increased in abundance by day 21 post-ALI, at which point the majority of the apical aspect of the culture was composed of ciliated cells. This increase in the number of ciliated cells following culturing for several weeks at ALI has similarly been observed in human AECMs [31, 32, 58]. There was no variation in the population of ciliated cells at day 42 post-ALI (Fig 4D and E). The ultrastructure of ciliated cells was further studied using SEM and TEM (Fig 5). The cilia present in the model were of correct morphology and comparable to ciliated cells present in *ex vivo* tissue, both in the structure of the basal bodies (Fig 5B) and 9 + 2 axoneme (Fig 5C). The length and density of the cilia also appears comparable (Fig 5A). Using light microscopy, it has been shown that cilia are capable of actively beating. Using microspheres, we have shown that this beating is in a co-ordinated fashion, demonstrating active mucociliary clearance on the apical aspect (data not shown).

The presence of mucus, secreted by goblet cells onto the apical surface, is typical of the airway epithelium [59]. Mucins ensnare invading particles, which are subsequently removed from the respiratory tract through mucociliary clearance [60]. Evidence of jacalin-labelling indicative of Muc5AC-positive goblet cells could be observed throughout the culturing of the BBECs, including during submerged culture (Fig 6A & S7 Fig) [28]. Mucus secretions from ALI-grown AECM have been confirmed to be highly representative to *in vivo* secretions [61], suggesting our model was capable of accurately replicating the mucosal phenotype of the bovine respiratory tract. The release of mucus at the apical surface was observed using SEM (Fig 6C). The presence of goblet cells actively extruding mucus into the apical compartment could be identified in the model between day 15-42 post-ALI (Fig 6C [iii]). Smaller globules of mucus were also present entrapped in cilia (Fig 6C [i]). This suggests that actively excreted mucus was only present in differentiated cultures.

Progenitor basal cells were identified in the model using the marker p63 [62]. At all time-points examined, basal cells could be seen present in the epithelium, forming a continuous row along the basal aspect (S3 Fig). This distribution reflected *ex vivo* tissue, in which a row of basal cells forms, attached to the basement membrane (Fig 2B). The number of basal cells remained constant, regardless of the differentiation state of the epithelium, as to be expected [63]. Basal cells act to repair and regenerate the epithelium following damage [64]. In our bovine AECM, we have been able to show regeneration of our model following injury using a scratch assay (data not shown). This suggests basal cells are capable of repairing the epithelium following damage to the bovine AECM.

Despite the high degree of similarity between the bovine respiratory tract epithelia and differentiated BBEC cultures, there were several differences. The BBEC cultures were consistently thinner than the epithelium of bovine bronchi, at both the distal and proximal positions (S1 Fig). Similarly, the lawn of cilia produced by the model had a lower degree of coverage in comparison to *ex vivo* tissue (S6 Fig). Variation between ALI cultures and their source tissue has also previously been reported in human AECMs [28]. This may be due to the BBEC cultures not reaching full differentiation as achieved *in vivo*, or alternatively due to a higher number of cells differentiating into goblet cells as opposed to ciliated cells in the model. This may be due to the process of de-differentiation and re-differentiation that has to occur during the culturing of the airway epithelial cells *in vitro*.

In this study we have fully characterised a differentiated AECM derived from bronchial epithelial cells isolated from cattle. This model was shown to be highly representative of the *ex vivo* epithelium from which the cell were derived. The degree of differentiation of the model was determined at three day intervals over a six-week period. The model was shown to be fully differentiated between day 21-42 post-ALI, providing a three week window during which the model is suitable for experimentation. The bovine AECM contained the major epithelial cell types of the bronchial epithelium (ciliated-, goblet- and basal cells) in a columnar, pseudostratified epithelium which was highly reflective of the *in vivo* epithelia. The hallmark defences of the respiratory tract, specifically barrier function and mucocilary clearance, were present, ensuring the model is an excellent mimic of the respiratory microenvironment. Use of BBECs provides a lost-cost, easily obtainable alternative to human AECM. The model is highly stable for extended periods of culturing and displays limited inter-donor variability. As such the bovine AECM described provides an excellent model of the bronchial epithelium for use in infection and toxicology experiments.

## Acknowledgements

We thank Ms Margaret Mullin and Ms Lynne Stevenson (both University of Glasgow) for assistance with electron microscopy and histology, respectively.

## Supplementary Figure Legends

**Supplementary Figure 1.**
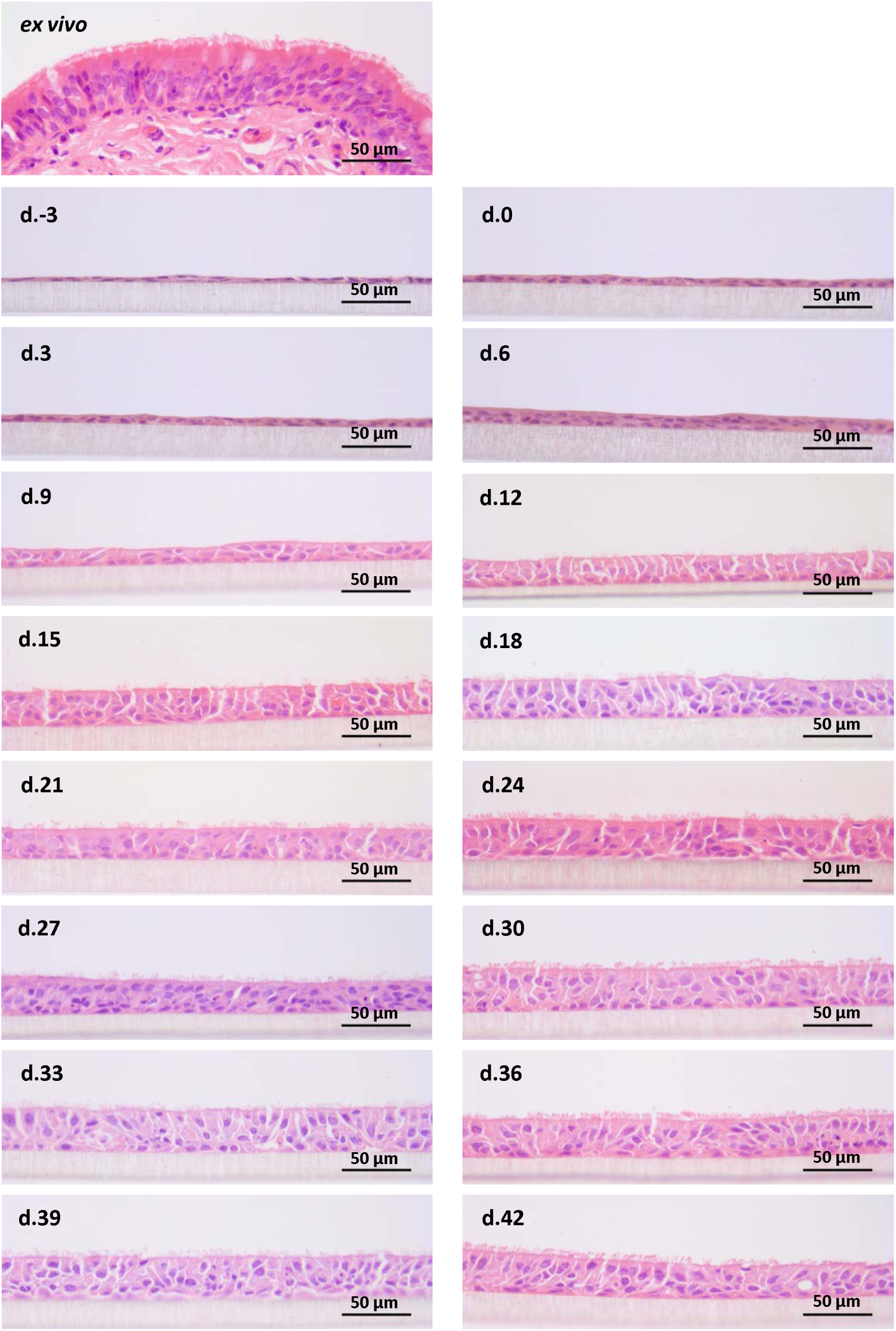
Histological assessment of epithelial morphology of BBEC cultures over time. BBEC cultures were grown for a stated number of days at an ALI before being fixed, paraffin-embedded and sections cut using standard histological techniques; sample of *ex vivo* tissue were also taken from each donor animal. Sections were subsequently deparaffinised and H&E stained.

**Supplementary Figure 2.**
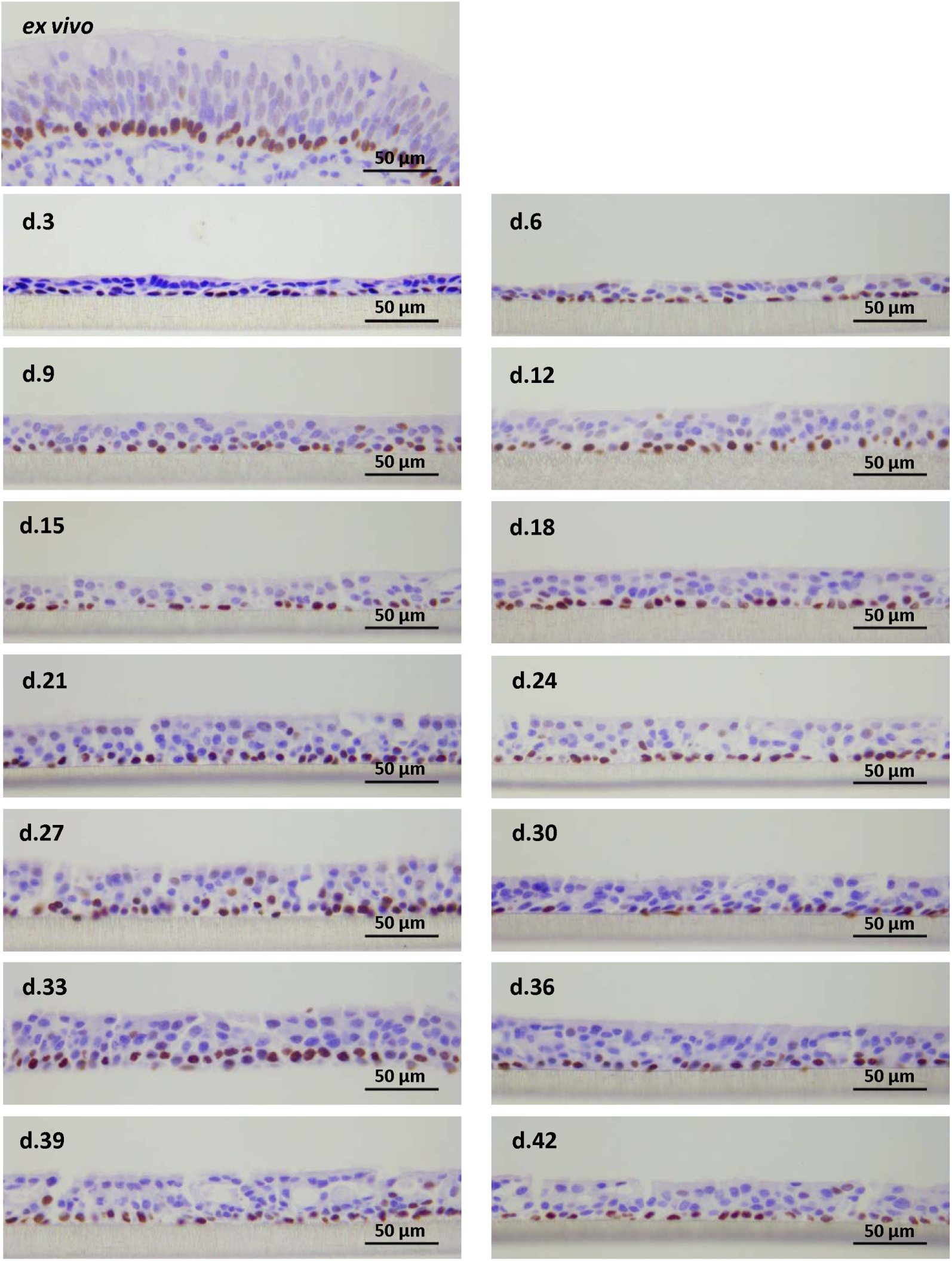
Histological assessment of basal cell distribution in BBEC cultures over time. BBEC cultures were grown for a stated number of days at an ALI before being fixed, paraffin-embedded and sections cut using standard histological techniques; sample of *ex vivo* tissue were also taken from each donor animal. Sections were subsequently deparaffinised and immunohistochemistry labelling of basal cells was performed using an anti-p63 antibody (positively labelled cells display brown-labelled nuclei). For days −3 and 0 the tissue layers were too thin to recover following antigen retrieval.

**Supplementary Figure 3.**
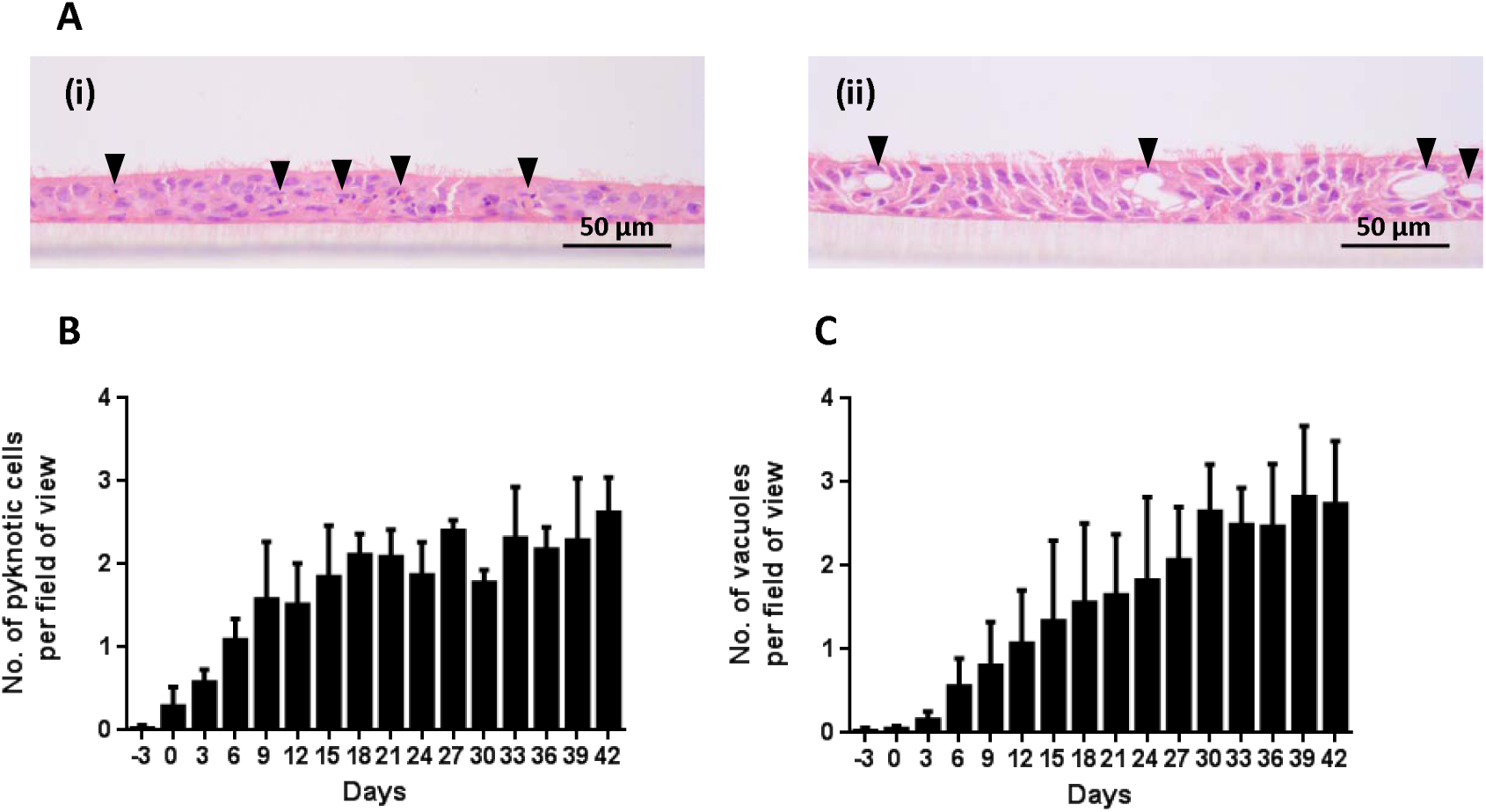
Assessment of deterioration in BBEC cultures over time from histological sections. BBEC cultures were grown for the stated number of days at an ALI before being fixed and paraffin-embedded using standard histological techniques. Sections were cut, deparaffinised and stained using H&E. Representative images in (A) are shown of (i) pyknotic cells (day 27 post-ALI) and (ii) epithelial gaps (day 33 post-ALI). Quantitative analysis (using ImageJ) of histological sections of BBEC layers fixed at three day intervals ranging from day −3 pre-ALI to day 42 post-ALI (see S1 Fig), showing (C) the number of pyknotic cells and (D) the number of epithelial gaps per field of view. For each insert, the numbers of pyknotic cells and epithelial gaps were counted in each of five 400x fields of view evenly distributed across the sample; three inserts were analysed per growth condition and the data represents the mean +/- standard deviation from tissue derived from three different animals.

**Supplementary Figure 4.**
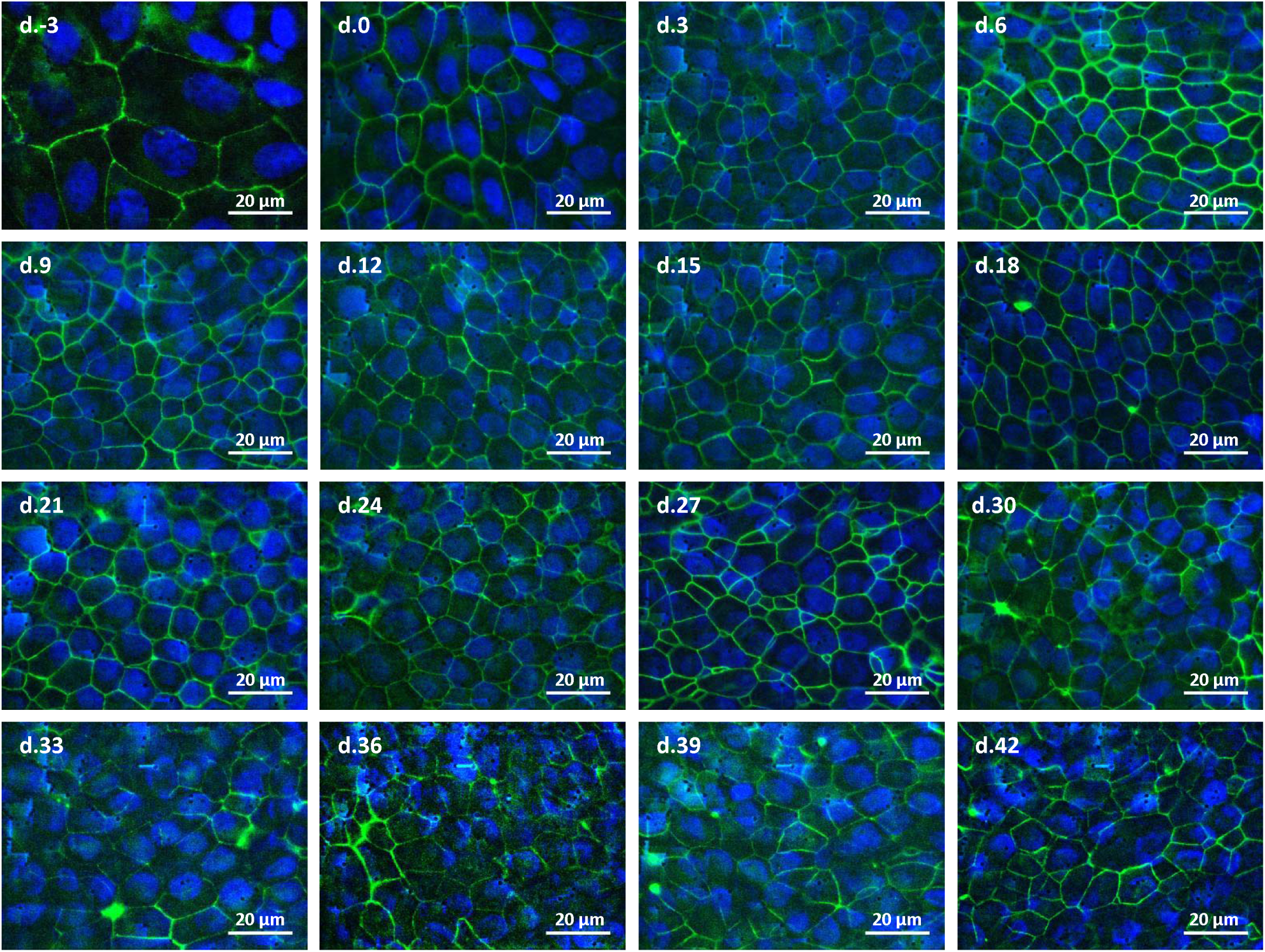
Tight junction formation in BBEC cultures over time assessed using immunofluorescence. BBEC cultures were grown for the stated number of days at an ALI before fixation. Samples were subsequently immunofluorescently stained for tight junctions (ZO-1 - green; nuclei - blue).

**Supplementary Figure 5.**
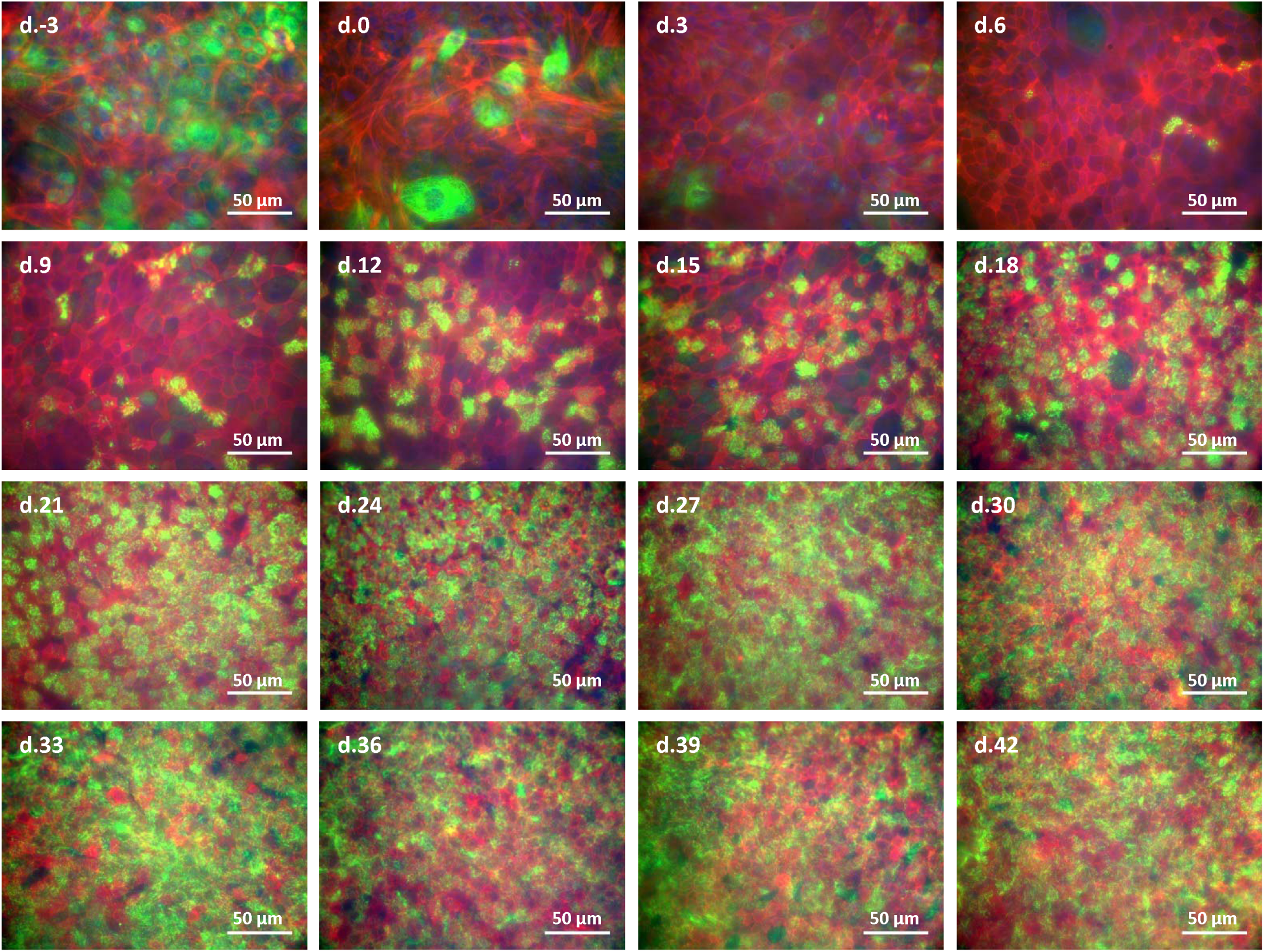
Differentiation of ciliated cells in BBEC cultures over time assessed using immunofluorescence. BBEC cultures were grown for the stated number of days at an ALI before fixation. Samples were subsequently immunofluorescently stained for cilia formation (β-tubulin - green; F-actin - red; nuclei - blue).

**Supplementary Figure 6.**
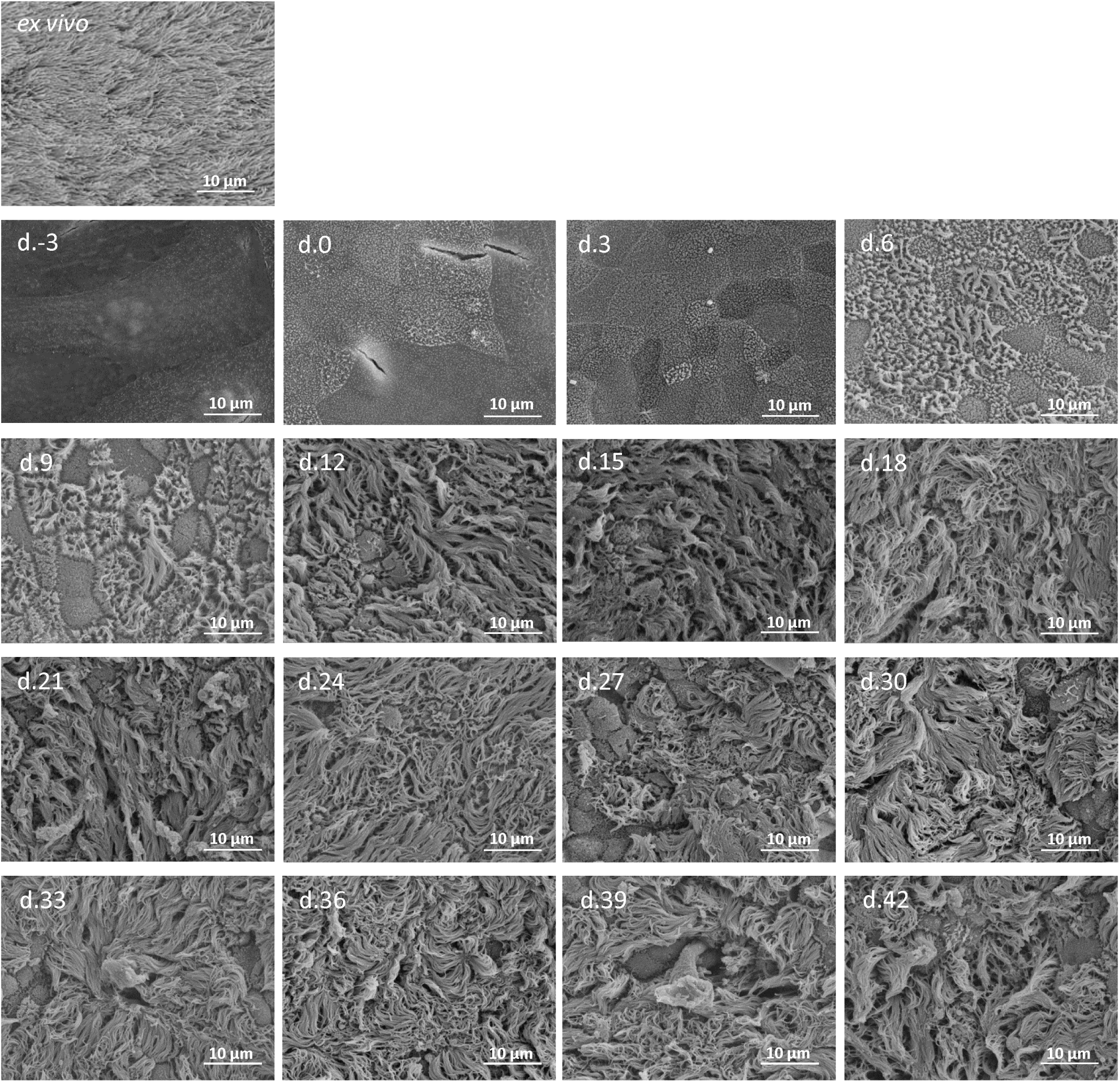
Differentiation of ciliated cells in BBEC cultures over time assessed using SEM. BBEC cultures were grown for the stated number of days at an ALI before fixation and processing for SEM. *Ex vivo* tissues were dissected prior to cell extraction and were also fixed, processed and analysed by SEM.

**Supplementary Figure 7.**
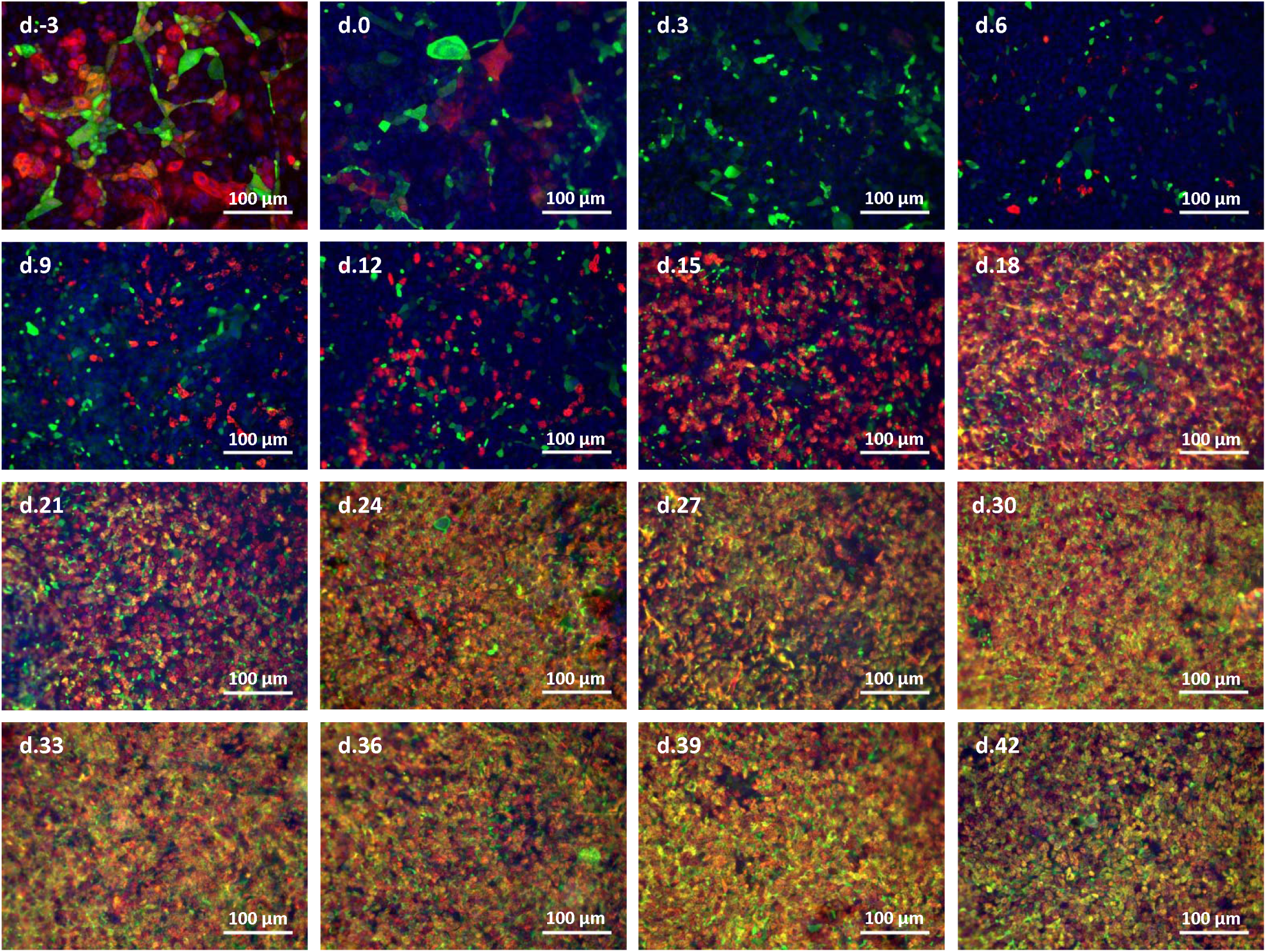
Differentiation of goblet cells in BBEC cultures over time assessed using immunofluorescence. BBEC cultures were grown for the stated number of days at an ALI before fixation. Samples were subsequently immunofluorescently stained for mucus-producing cells (Muc5AC - green; β-tubulin - red; nuclei - blue).

